# Precision Identification of Diverse Bloodstream Pathogens from the Gut Microbiome

**DOI:** 10.1101/310441

**Authors:** Fiona B. Tamburini, Tessa M. Andermann, Ekaterina Tkatchenko, Fiona Senchyna, Niaz Banaei, Ami S. Bhatt

## Abstract

Bloodstream infection is the most common infectious complication in hematopoietic cell transplantation recipients. To evaluate the genomic concordance of bloodstream pathogens and bacterial strains within the intestinal microbiome using whole genome sequencing, we developed StrainSifter, a bioinformatic pipeline to compare nucleotide variation between bacterial isolate strains and stool metagenomes. We applied StrainSifter to bloodstream isolates and stool metagenome samples from hematopoietic stem cell transplant recipients with bloodstream infections. StrainSifter is designed to identify single nucleotide variants between isolate and metagenomic short reads using stringent alignment, coverage, and variant frequency criteria for strain comparison. We identified enteric BSI isolates that were highly concordant with those in the gut microbiota, as well as highly concordant strains of typically non-enteric bacteria. These findings demonstrate the utility of StrainSifter in strain matching and provide a more precise investigation of the intestine as a reservoir of diverse pathogens capable of causing bloodstream infections.

## Introduction

Microbial pathogenicity and transmission depend in part on strain-level variability in both gene content and at the single-nucleotide variant (SNV) level, as different strains of the same organism can vary widely in their ability to cause disease (Snitkin et al. 2011; Lieberman et al. 2013). Whole genome sequencing (WGS) has facilitated the exploration of strain-level determinants of virulence and has enabled the precise tracking of pathogen transmission (Snitkin et al. 2011; Snitkin et al. 2012). Methods for strain tracking and comparison most commonly rely on either sequencing a defined number of genes or identifying single nucleotide variants (SNVs) across the whole genome. For example, whole genome sequencing of cultured *Klebsiella pneumoniae* stool isolates in ICU patients demonstrated the role of *K. pneumoniae* colonization in systemic infections (Gorrie et al. 2017; Martin et al. 2016; Nielsen et al. 2017) and genome-wide SNV comparison is routinely employed for epidemiologic strain tracking (Salipante et al. 2015; Snitkin et al. 2012; Harris et al. 2013; Chen et al. 2017; Kovanen et al. 2014). While strain comparisons have primarily been performed on bacterial isolates, newer computational tools (MIDAS, StrainPhlAn, and metaSNV) profile strain variation between metagenomic samples over time without the bias inherent in culturing isolates (Costea et al. 2017; Nayfach et al. 2016; Truong et al. 2017).

Despite the increasing use of WGS for strain comparison, bioinformatic tools have not been developed for the specific purpose of comparing disease-causing bacterial isolates to metagenomic samples at the nucleotide level within the same patient. It is desirable to achieve the highest resolution of comparison across the whole genome of an organism in order to precisely compare clinical isolates to strains within a sampled metagenome. Additionally, most tools depend on the availability of reference genomes and are thus often restricted to studying well characterized bacterial pathogens, limiting the types of organisms that are available for strain comparisons. To date, no reference-free tool exists with the stringent filtering steps necessary to identify genomic variants specific to a single strain between genomes and metagenomes. We sought to compare strain-level identity between bloodstream infection isolates and stool metagenomes using samples from immunocompromised patients at elevated risk of systemic infections, with the goal of better understanding the origins of bloodstream infection in this patient population.

Hematopoietic stem cell transplant (HCT) recipients are at high risk for bloodstream infections, which occur in 20-60% of patients primarily during periods of neutropenia and mucositis (Blennow et al. 2014; Gudiol et al. 2014; Mikulska et al. 2009). Prior to engraftment of donor stem cells, HCT recipients are particularly susceptible to translocation of enteric organisms from the intestinal microbial reservoir into the bloodstream. During periods of mucositis, primary bloodstream infections with enteric Gram-negative and Gram-positive organisms are thought to arise from mucosal barrier injury (See et al. 2017). In contrast, non-enteric commensal and environmental bacteria access the bloodstream through other routes including intravenous lines and sites where skin epithelial integrity has become compromised. Existing methods for strain comparison have been helpful in identifying the origins of bloodstream infections in HCT patients using culture-based pulsed-field gel electrophoresis (PFGE), or PCR-based multi-locus sequence typing (MLST) (Tancrede and Andremont 1985; Samet et al. 2013). Although these methods are rapid, affordable, and standardized across many organisms, they have several limitations: 1) they require isolation and culture of candidate pathogens from the stool or other suspected pathogen reservoirs, a step that can introduce selection bias, and 2) they often rely on detailed characterization of a small number of genes and therefore lack the discriminatory power required for robust, genome-wide strain comparison (W. Li, Raoult, and Fournier 2009). No studies to date have investigated the origins of bloodstream infections in HCT or other immunocompromised patients at the level of single nucleotide variation across entire genomes.

In this work, we present StrainSifter, a straightforward bioinformatics pipeline based on whole genome sequencing to comprehensively and precisely compare single nucleotide variants between isolate genomes and metagenomes without bias introduced by culturing. Additionally, unlike most previous strain comparison bioinformatic tools (MIDAS and StrainPhlAn), variant calling within our pipeline does not depend on the availability of a reference genome database, a particularly relevant consideration in immunocompromised patients who develop infections with lesser known and poorly studied opportunistic pathogens. We have applied this tool to track bacterial strain translocation from the gut to the bloodstream in a unique cohort of immunocompromised patients at high risk for translocation and bloodstream infection, specifically patients with hematologic malignancies who have undergone hematopoietic stem cell transplantation. We find that StrainSifter is able to identify strains of both intestinal mucosa-associated *(Enterococcus faecium, Escherichia coli, Klebsiella pneumoniae, Enterobacter cloacae, and Streptococcus mitis)* and typically non-mucosal bacteria *(Staphylococcus epidermidis* and *Pseudomonas aeruginosa*) within the gut microbiome as being concordant with the same strain cultured from the blood of patients with bloodstream infections. Intriguingly, this suggests that both enteric and classically nonenteric pathogens can colonize and even thrive within the intestinal reservoir of immunocompromised hosts.

## Methods

### Cohort selection

A retrospective cohort study, approved by the institutional review board under the IRB protocol #8903 (PI: Dr. David Miklos, co-PI: Dr. Ami Bhatt), was performed at Stanford Hospital. Informed consent was obtained from all subjects prior to specimen collection. At the time of cohort identification (July 2017), a stool biospecimen collection containing 964 stool samples from 402 patients was available for investigation. This collection consisted of convenience samples collected from autologous and allogeneic stem cell transplantation patients at Stanford University Hospital between October 5 2015 and June 9 2017. Patients were included in this study if a stool sample had been collected within 30 days prior to an episode of bloodstream infection (BSI) for which a blood isolate was also available. From this final cohort, we sequenced all stool samples in our collection within 60 days prior to and 30 days after BSI.

### Bloodstream isolate identification

Bloodstream isolates from HCT patients who received medical care at Stanford University Hospital were obtained from the Stanford Hospital Clinical Microbiology Laboratory. All isolates considered typical bloodstream pathogens by National Healthcare Safety Network (NHSN) guidelines are stored in a glycerol suspension at -80°C for 12 months (“Centers for Disease Control and Prevention” 2018). Blood culture isolates considered to be skin-associated bacteria (including viridans group *Streptococcus spp.* and coagulase-negative *Staphylococcus spp*.) were saved if they were recovered in two or more blood culture sets as per NHSN criteria (“Centers for Disease Control and Prevention” 2018). Isolates were identified by standard biochemical testing and matrix-assisted laser desorption and ionization time-of-flight mass spectrometry (MALDI-TOF) (Bruker Daltonics, Bremen, Germany).

### Sample processing

Bloodstream bacterial isolates were plated on brain heart infusion agar with 10% horse blood. DNA was extracted from isolates using the Gentra Puregene Yeast/Bact. Kit per manufacturer’s instructions. Stool samples were collected and stored at 4°C for up to 24 hours prior to homogenization, aliquoting and storage at 80°C. DNA was extracted from stool using the QIAamp DNA Stool Mini Kit (QIAGEN^®^) per manufacturer’s instructions, with an initial bead-beating step prior to extraction using the Mini-Beadbeater-16 (BioSpec Products) and 1 mm diameter Zirconia/Silica beads (BioSpec Products). Bead-beating consisted of 7 rounds of alternating 30 second bead-beating bursts followed by 30 seconds cooling on ice. DNA concentration for all samples was measured using Qubit^®^ Fluorometric Quantitation (Life Technologies). DNA sequencing libraries from both isolates and stool were prepared using the Nextera XT DNA Library Prep Kit (Illumina^®^) with isolates and stool microbiota libraries prepared at separate times following DNAse treatment of all lab surfaces to avoid cross-contamination. Library concentration was measured using Qubit^®^ Fluorometric Quantitation (Life Technologies) and library quality and size distributions were analyzed with the Bioanalyzer 2100 (Agilent). Prepared libraries were multiplexed and subjected to 100 base pair, paired-end sequencing on the HiSeq 4000 platform (Illumina^®^).

### Shotgun sequencing analysis

#### Preprocessing

Sequencing data were demultiplexed by unique barcodes (bcl2fastq v2.20.0.422, Illumina^®^). Read quality was assessed using FastQC v0.11.4. Reads were deduplicated to remove PCR and optical duplicates using SuperDeduper v1.4 with the start location in the read at 5 bp (-s 5) and minimum length of 50 bp (-l 50). Deduplicated reads were trimmed using TrimGalore v0.4.4, a wrapper for CutAdapt v1.16, with a minimum quality score of 30 for trimming (-q 30), minimum read length of 50 (--length 50) and the “--nextera” flag to remove Illumina Nextera adapter sequences. Draft genomes of bacterial isolates were assembled using SPAdes v3.8.0 (Bankevich et al. 2012) with default parameters. Summary statistics for each BSI assembly were generated using ‘basic_assembly_stats.py’ from GAEMR v1.0.1 (“GAEMR” n.d).

#### Taxonomic classification

Bloodstream isolates were taxonomically classified by aligning draft genome contigs to the BLAST nt database and identifying the most common top hit for each contig. Gut metagenomic reads were taxonomically classified via the One Codex platform, a web-based tool for assigning read-level classifications based on unique k-mer signatures relative to a curated reference database [2017 database] (Minot, Krumm, and Greenfield 2015).

#### Phylogenetic tree building and variant identification with the StrainSifter pipeline

StrainSifter is a pipeline deployed via Snakemake with modules for variant calling and phylogenetic tree building, available at GitHub (https://github.com/bhattlab/strainsifter). StrainSifter accepts as input an assembled bacterial draft genome and two or more short read data sets (isolate or metagenomic), and can report a phylogenetic tree of input samples as well as pairwise SNV counts. To build the phylogenetic trees reported in this manuscript, an NCBI RefSeq reference genome for each infectious species (based on clinical laboratory classifications) was provided to StrainSifter. For the variant counting reported herein, BSI draft genomes were supplied to StrainSifter. For both analyses, all stool and BSI short read datasets were provided as input. For both phylogeny and SNV-counting modules, preprocessed short reads are first aligned to the reference genome using the Burrows-Wheeler aligner (BWA) v0.7.10 (Li and Durbin 2009). Alignments are filtered to include only high-confidence alignments with mapping quality of at least 60 using the “view” tool from the SAMtools suite (v1.7)(Li et al. 2009) (samtools view -b -q 60), and further filtered using BamTools “filter” (v2.4.0) to include only reads with the desired number or fewer mismatches (i.e. for five or fewer mismatches: bamtools filter -tag ‘NM:<6’) (Barnett et al. 2011) For phylogenetic tree construction, reads with 5 or fewer mismatches were included; for determining strain single-nucleotide variants, reads were limited to 1 or fewer mismatches. Per-base coverage is calculated from each resulting BAM file using bedtools genomecov (v2.26.0) and processed with custom python scripts to identify samples meeting a minimum average coverage of 5X across at least 50% of the genome. Only samples meeting the coverage requirement are continued through the pipeline. Pileup files are created from BAM files using SAMtools “mpileup”, and are analyzed using custom python scripts to identify bases occurring with at least 0.8 frequency at positions covered 5X or greater. Only bases with a minimum phred score of >=20 are considered. Consensus sequences for each sample are created, wherein bases that cannot be confidently determined given the described parameters are called as “N”.

To create a phylogenetic tree, core positions are identified on a per-species basis, where core positions are defined as positions in the reference genome where a base could be confidently called for all samples analyzed. To generate phylogenetic trees, core positions with variants in at least one sample are identified and concatenated into fasta files. Fasta files are aligned using MUSCLE v3.8.31 (Edgar 2004) and a maximum-likelihood phylogenetic tree is computed using FastTree v2.1.7 (Price, Dehal, and Arkin 2010). Phylogenetic trees are visualized in R using the ape, phangorn, and ggtree packages. Pairwise SNVs are determined from the consensus sequences using a custom python script.

#### Synthetic multilocus sequence typing

Metagenomic short reads were assembled using metaSPAdes v3.8.0 (Nurk et al. 2017). Multilocus sequence typing (MLST) schemes and sequences were downloaded from the PubMLST database (“PubMLST” n.d.). MLST gene sequences were aligned to metagenome assemblies, and the top hit for each alignment was chosen based on E-value, percent identity, and alignment length. Only MLST sequences that were present in the metagenomic assembly with 100% identity across the entire length of the sequence were reported. MLST types generated by our in-house analysis were confirmed with the SRST2 synthetic multilocus sequence typing tool (Inouye et al. 2014).

#### Plots

Plots were generated using the R programming language using the ggplot2, reshape2, and dplyr packages.

## Results

### Sample collection and cohort

During the study period, we identified 74 patients within our stool biospecimen collection with a bloodstream infection (77 BSI episodes). Of these, we identified 30 patients (32 bloodstream isolates) with at least 1 collected stool sample within 30 days prior to their bacterial BSI. From our final cohort of 30, we sequenced all stool samples collected between 60 days prior to and 30 days after the date of BSI, 83 samples in total. Patients had a median of 2 stool samples (range 1-8), with the sampling distribution presented in Figure 1. Stools were collected at a median of 9 days prior to BSI (range -58 to +31). Clinical characteristics of the 30 patients are listed in Table 1 (see Table S1 for individual patient data). All patients in the cohort were on at least one antibiotic within 30 days prior to BSI, with fluoroquinolones being the most commonly administered antibiotic.

**Figure 1.**
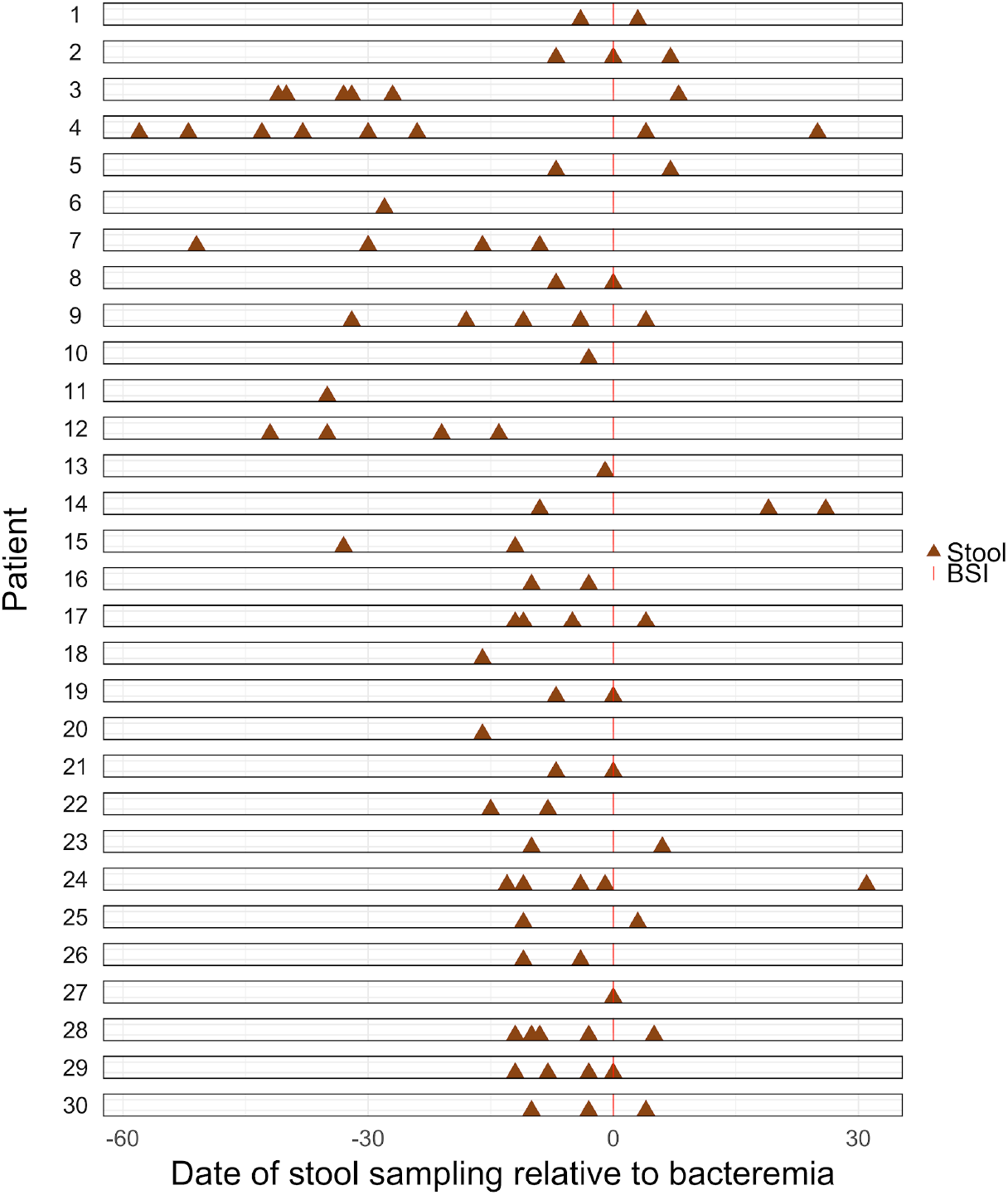
Stool sampling. Timeline of stool sampling relative to BSI and hematopoietic cell transplant for patients included in the study.

**Table 1.**
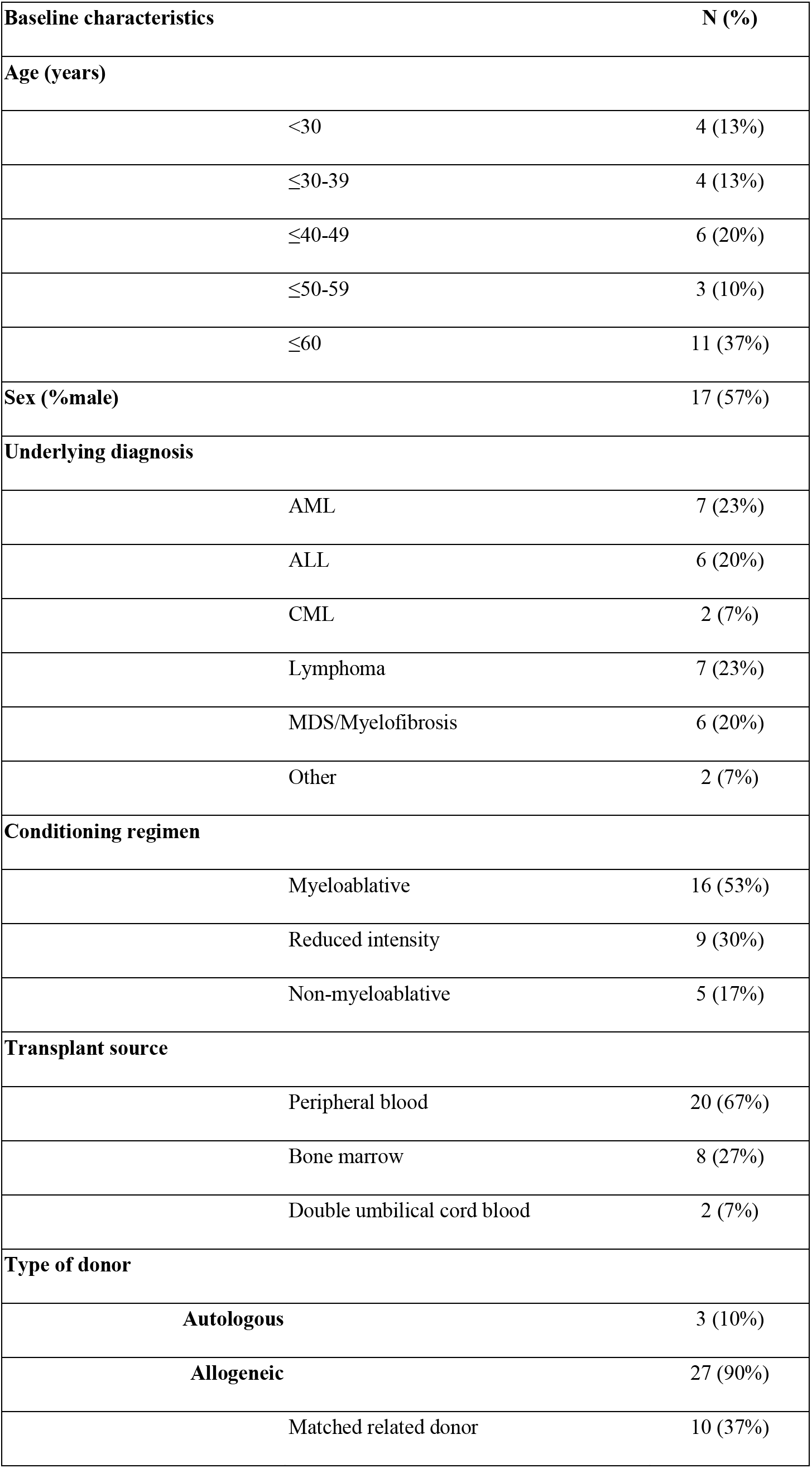
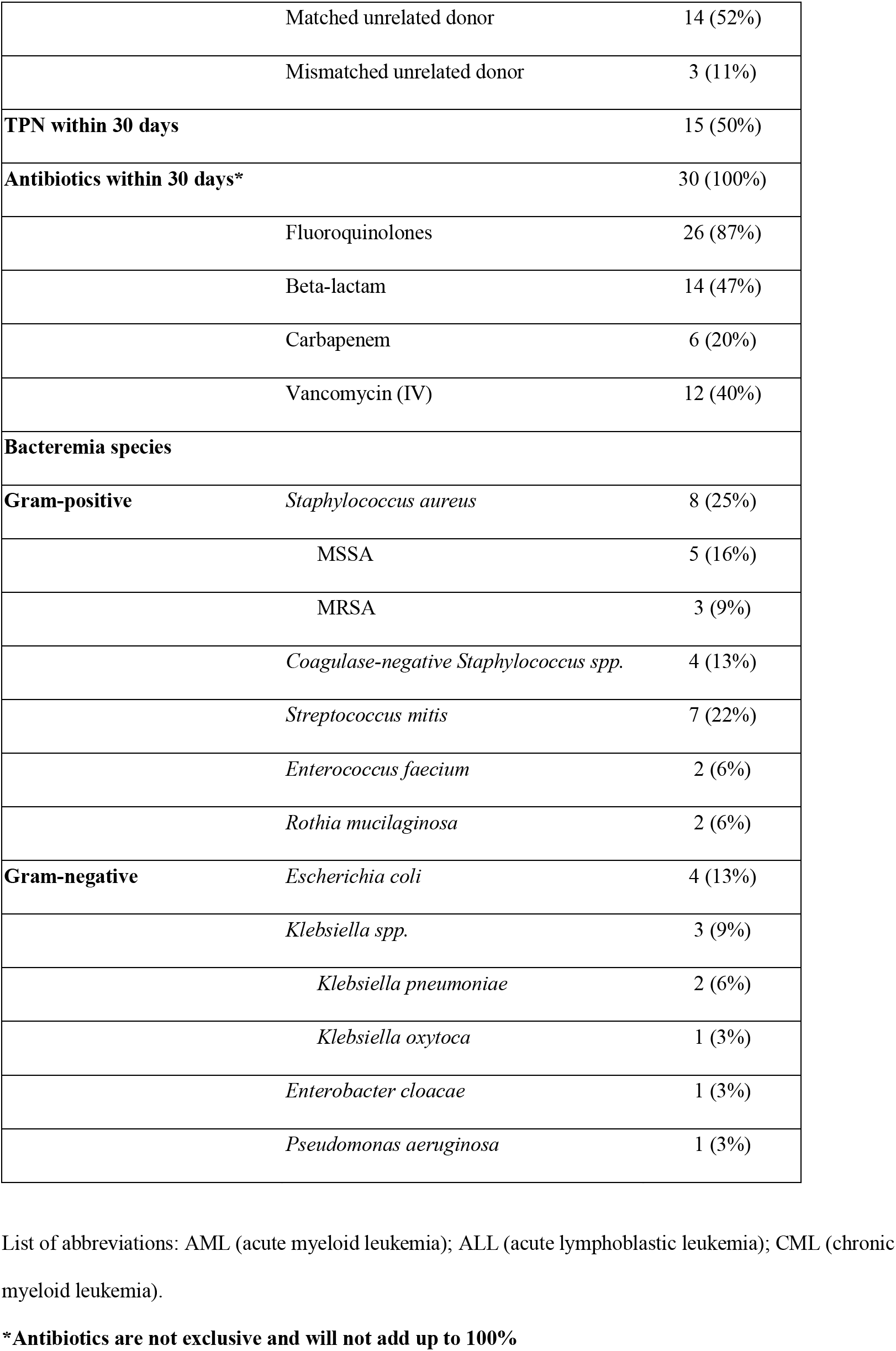
List of abbreviations: AML (acute myeloid leukemia); ALL (acute lymphoblastic leukemia); CML (chronic myeloid leukemia). **\*Antibiotics are not exclusive and will not add up to 100%**

| **Baseline characteristics** | **N (%)** |
| --- | --- |
| **Age (years)** |  |
| <30 | 4 (13%) |
| ≤30-39 | 4 (13%) |
| ≤40-49 | 6 (20%) |
| ≤50-59 | 3 (10%) |
| ≤60 | 11 (37%) |
| **Sex (%male)** | 17 (57%) |
| **Underlying diagnosis** |  |
| AML | 7 (23%) |
| ALL | 6 (20%) |
| CML | 2 (7%) |
| Lymphoma | 7 (23%) |
| MDS/Myelofibrosis | 6 (20%) |
| Other | 2 (7%) |
| **Conditioning regimen** |  |
| Myeloablative | 16 (53%) |
| Reduced intensity | 9 (30%) |
| Non-myeloablative | 5 (17%) |
| **Transplant source** |  |
| Peripheral blood | 20 (67%) |
| Bone marrow | 8 (27%) |
| Double umbilical cord blood | 2 (7%) |
| **Type of donor** |  |
| **Autologous** | 3 (10%) |
| **Allogeneic** | 27 (90%) |
| Matched related donor | 10 (37%) |

|  |  |  |
| --- | --- | --- |
|  | Matched unrelated donor | 14 (52%) |
|  | Mismatched unrelated donor | 3 (11%) |
| **TPN within 30 days** |  | 15 (50%) |
| **Antibiotics within 30 days\*** |  | 30 (100%) |
|  | Fluoroquinolones | 26 (87%) |
|  | Beta-lactam | 14 (47%) |
|  | Carbapenem | 6 (20%) |
|  | Vancomycin (IV) | 12 (40%) |
| **Bacteremia species** |  |  |
| **Gram-positive** | *Staphylococcus aureus* | 8 (25%) |
|  | MSSA | 5 (16%) |
|  | MRSA | 3 (9%) |
|  | *Coagulase-negative Staphylococcus spp.* | 4 (13%) |
|  | *Streptococcus mitis* | 7 (22%) |
|  | *Enterococcus faecium* | 2 (6%) |
|  | *Rothia mucilaginosa* | 2 (6%) |
| **Gram-negative** | *Escherichia coli* | 4 (13%) |
|  | *Klebsiella spp.* | 3 (9%) |
|  | *Klebsiella pneumoniae* | 2 (6%) |
|  | *Klebsiella oxytoca* | 1 (3%) |
|  | *Enterobacter cloacae* | 1 (3%) |
|  | *Pseudomonas aeruginosa* | 1 (3%) |

### Infectious taxa are present in the gut microbiome near the time of infection

We sequenced an average of 68,704,214 reads per stool sample (20,298,792-148,162,892) and 27,999,060 reads per isolate (10,652,310-61,058,906) (Table S2). Isolate reads were assembled into draft genomes (Table S3).

MALDI-TOF is unable to discriminate between some closely related taxa (e.g. *Streptococcus* spp.), so we sought to confirm the clinical laboratory BSI classifications by aligning draft genome contigs to the

BLAST nt database. Of 32 BSIs, we identified 3 for which WGS and MALDI-TOF results disagreed. Two *Streptococcus* isolates that had been classified as *Streptococcus mitis* by MALDI-TOF were identified as *Streptococcus oralis,* and one as *Streptococcus pseudopneumoniae* by WGS (Table S4).

To determine community composition of pre- and post-BSI stool samples, we taxonomically classified gut metagenomic reads via the One Codex platform. Considering species level read classifications, we observe the target species in the gut at a threshold of 0.1% or greater relative abundance for 19 of 83 stools (23%) or 20 of 95 stool-BSI pairs (21%) taking into account episodes of polymicrobial BSI; one patient developed a BSI with two bacterial species, both of which were present in the stool above threshold level (Table 2; full taxonomic classifications in table S5). Read-level taxonomic classifiers are often unable to discriminate between closely related species, so we also compared BSIs and stool read classifications at the genus level. We observe the target genus in the gut community for 27% of stools, or 23% of stool-BSI pairs (Table S5, S6).

**Table 2.**

| Patient | Day relative to BSI | BSI organism | Relative abundance (%) |
| --- | --- | --- | --- |
| 1 | -4 | Staphylococcus aureus | 0 |
| 1 | 3 | Staphylococcus aureus | 0 |
| 2 | -7 | Enterobacter cloacae | 0 |
| 2 | 0 | Enterobacter cloacae | 0.74 |
| 2 | 7 | Enterobacter cloacae | 0 |
| 2 | -7 | Klebsiella pneumoniae | 0 |
| 2 | 0 | Klebsiella pneumoniae | 3.47 |
| 2 | 7 | Klebsiella pneumoniae | 0.01 |
| 3 | -27 | Escherichia coli | 87.15 |
| 3 | -32 | Escherichia coli | 97.25 |
| 3 | -33 | Escherichia coli | 96.4 |
| 3 | -40 | Escherichia coli | 0.05 |
| 3 | -41 | Escherichia coli | 0.43 |
| 3 | 8 | Escherichia coli | 0.01 |
| 3 | -27 | Staphylococcus aureus | 0 |
| 3 | -32 | Staphylococcus aureus | 0 |
| 3 | -33 | Staphylococcus aureus | 0 |
| 3 | -40 | Staphylococcus aureus | 0 |
| 3 | -41 | Staphylococcus aureus | 0 |
| 3 | 8 | Staphylococcus aureus | 0 |
| 4 | -24 | Staphylococcus aureus | 0 |
| 4 | -30 | Staphylococcus aureus | 0 |
| 4 | -38 | Staphylococcus aureus | 0 |
| 4 | -43 | Staphylococcus aureus | 0 |
| 4 | -52 | Staphylococcus aureus | 0 |
| 4 | -58 | Staphylococcus aureus | 0 |
| 4 | 25 | Staphylococcus aureus | 0 |
| 4 | 4 | Staphylococcus aureus | 0 |
| 5 | -7 | Staphylococcus aureus | 0 |
| 5 | 7 | Staphylococcus aureus | 0 |
| 6 | -28 | Klebsiella oxytoca | 0 |
| 7 | -16 | Escherichia coli | 0.79 |
| 7 | -30 | Escherichia coli | 0.22 |
| 7 | -51 | Escherichia coli | 2.05 |
| 7 | -9 | Escherichia coli | 0.38 |
| 8 | -7 | Rothia mucilaginosa | 0 |
| 8 | 0 | Rothia mucilaginosa | 0 |
| 9 | -11 | Escherichia coli | 0.31 |
| 9 | -18 | Escherichia coli | 0 |
| 9 | -32 | Escherichia coli | 0 |
| 9 | -4 | Escherichia coli | 0.03 |
| 9 | 4 | Escherichia coli | 0 |
| 10 | -3 | Staphylococcus aureus | 0 |
| 11 | -35 | Escherichia coli | 0.04 |
| 12 | -14 | Staphylococcus aureus | 0 |
| 12 | -21 | Staphylococcus aureus | 0 |
| 12 | -35 | Staphylococcus aureus | 0.65 |
| 12 | -42 | Staphylococcus aureus | 0.09 |
| 13 | -1 | Staphylococcus epidermidis | 2.51 |
| 14 | -9 | Klebsiella pneumoniae | 0 |
| 14 | 19 | Klebsiella pneumoniae | 0 |
| 14 | 26 | Klebsiella pneumoniae | 0 |
| 15 | -12 | Staphylococcus epidermidis | 0 |
| 15 | -12 | Staphylococcus epidermidis | 0 |
| 15 | -33 | Staphylococcus epidermidis | 0 |
| 15 | -33 | Staphylococcus epidermidis | 0 |
| 16 | -10 | Streptococcus mitis | 0 |
| 16 | -3 | Streptococcus mitis | 0 |
| 17 | -11 | Streptococcus mitis | 0 |
| 17 | -12 | Streptococcus mitis | 0 |
| 17 | -5 | Streptococcus mitis | 0 |
| 17 | 4 | Streptococcus mitis | 0.03 |
| 18 | -16 | Staphylococcus epidermidis | 0 |
| 18 | -16 | Staphylococcus epidermidis | 0 |
| 19 | -7 | Pseudomonas aeruginosa | 0 |
| 19 | 0 | Pseudomonas aeruginosa | 0.83 |
| 20 | -16 | Staphylococcus epidermidis | 0 |
| 21 | -7 | Staphylococcus aureus | 0.01 |
| 21 | 0 | Staphylococcus aureus | 0.25 |
| 22 | -15 | Streptococcus mitis | 0 |
| 22 | -8 | Streptococcus mitis | 2.33 |
| 23 | -10 | Enterococcus faecium | 11.61 |
| 23 | 6 | Enterococcus faecium | 0 |
| 24 | -1 | Rothia mucilaginosa | 0.04 |
| 24 | -11 | Rothia mucilaginosa | 0 |
| 24 | -13 | Rothia mucilaginosa | 0 |
| 24 | -4 | Rothia mucilaginosa | 0 |
| 24 | 31 | Rothia mucilaginosa | 0 |
| 25 | -11 | Enterococcus faecium | 0.36 |
| 25 | 3 | Enterococcus faecium | 94.09 |
| 26 | -11 | Streptococcus mitis | 0 |
| 26 | -4 | Streptococcus mitis | 0 |
| 27 | 0 | Staphylococcus aureus | 0.51 |
| 28 | -10 | Streptococcus mitis | 0 |
| 28 | -12 | Streptococcus mitis | 0.04 |
| 28 | -3 | Streptococcus mitis | 0 |
| 28 | -9 | Streptococcus mitis | 0.02 |
| 28 | 5 | Streptococcus mitis | 0 |
| 29 | -12 | Streptococcus mitis | 0 |
| 29 | -3 | Streptococcus mitis | 0 |
| 29 | -8 | Streptococcus mitis | 0.01 |
| 29 | 0 | Streptococcus mitis | 0.03 |
| 30 | -10 | Streptococcus mitis | 0 |
| 30 | -3 | Streptococcus mitis | 0 |
| 30 | 4 | Streptococcus mitis | 0 |

### Gut domination with an infectious pathogen does not always precede bloodstream infection

We sought to determine whether bloodstream infection is preceded by gut microbiota domination by a pathogen as has been previously reported (Ubeda et al. 2010; Taur et al. 2012). We defined domination as 30% species relative abundance or greater of an organism in the gut, as previously described (Taur et al. 2012). Of the 19 stools in which the BSI organism was detected, only 4 stool samples (21%) were dominated by the BSI pathogen. Stool samples from patient 3 were dominated by *Escherichia coli* (>85% relative abundance) 27, 32, and 33 days prior to the development of an *E. coli* BSI (Figure 2A; table 2); *Enterococcus faecium* dominated the intestinal microbiome (94% relative abundance) in patient 25 three days after *E. faecium* BSI (Figure 2B; table 2).

**Figure 2.**
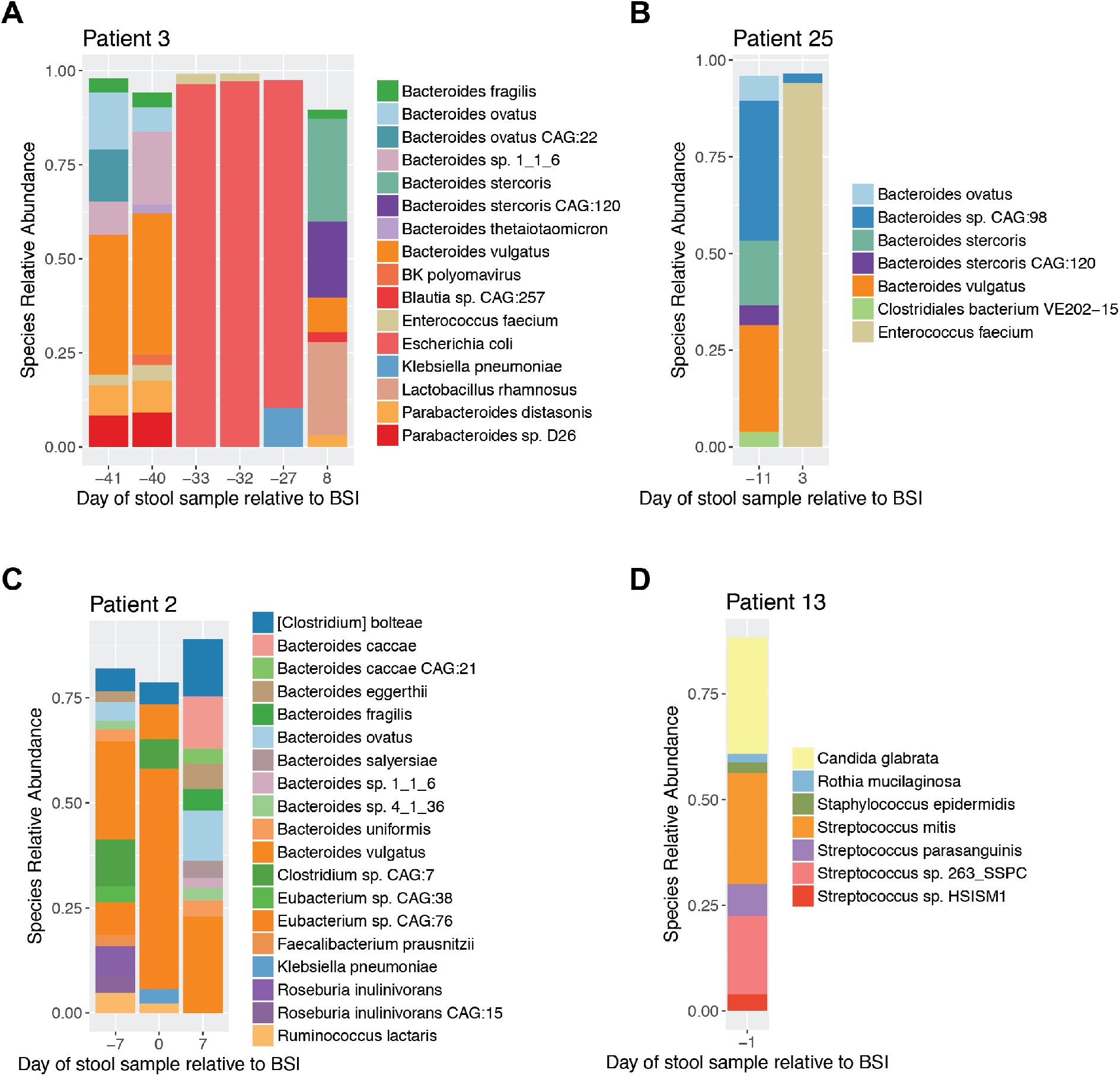
Gut domination by a pathogen does not always precede bloodstream infection. Domination by *E. coli* (A) and Enterococcus faecium (B) occurs prior to bacteremia, while *K. pneumoniae* (C) and *S. epidermidis* (D) are present in the gut microbiota of their respective patients prior to BSI but not at high abundance. Plots show species present at 2% relative abundance or greater (host reads excluded).

Several patients experienced BSI with organisms present at relatively lower abundance in the gut prior to bloodstream infection: *Staphylococcus epidermidis* at approximately 2.5% relative abundance in the gut of patient 13 one day prior to their *S. epidermidis* infection, *Pseudomonas aeruginosa* at low abundance (0.83%) in the gut of patient 19 on the day of infection, and *Klebsiella pneumoniae* at 3.5% relative abundance in the gut of patient 2, 9 days prior to BSI (Figure 2C,D; table 2).

We also observe gut domination by non-BSI-causing pathogens. Patient 14 experienced a *K. pneumoniae* BSI, yet stool samples at two timepoints were dominated by other pathogens: *E. coli* at 67% relative abundance 9 days prior to BSI and *E. faecium* at 85% relative abundance 19 days after BSI (Table 2).

### Phylogenetic analysis indicates close relatedness of gut and BSI strains

We developed the StrainSifter pipeline to perform nucleotide-level comparisons between genomes and metagenomes. StrainSifter is deployed as a Snakemake workflow, and the pipeline steps are described in Figure 3. Briefly, the inputs to StrainSifter are a draft genome (the “target” genome) and one or more other short read datasets (isolate genomes or metagenomes). StrainSifter detects whether the target organism is present with sufficient abundance in the short read datasets and outputs phylogenetic trees and SNV counts between samples. Despite an abundance of research studies focused on strain-level comparisons of microbes, there is still no clear definition of what constitutes a “strain,” and it is unclear whether genomes that differ by a few nucleotides should be considered distinct strains. To investigate the overall relatedness of strains of each BSI species in our metagenomes and isolates, we first took a comparative phylogenomics approach by applying StrainSifter to assess inter-patient strain relatedness. Rather than compare strains from the same patient to publicly-available reference genomes, we instead compared the phylogenetic relatedness of a patient’s gut and BSI strains to strains from the other patient samples in our collection. We reasoned that strains from the same geographic location are more likely to be closely phylogenetically related than strains that are both spatially and temporally distant, so comparing inter-patient strain relatedness in the context of the other strains in our sample collection should more precisely indicate strains or lineages unique to a patient.

**Figure 3.**
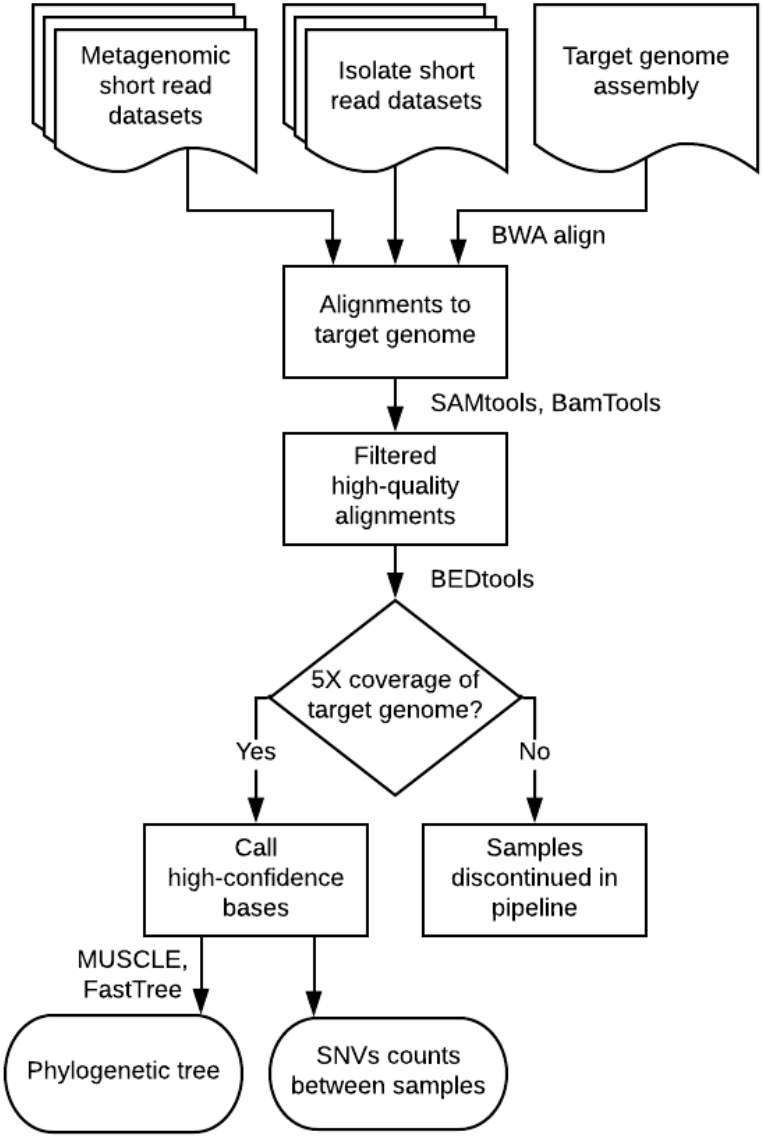
StrainSifter pipeline. Flowchart showing input, analysis steps, and output of the StrainSifter pipeline.

For each BSI species, we used StrainSifter to identify metagenomic samples with at least 5X coverage of the NCBI representative genome for each target organism, found core SNVs present in each sample, and inferred maximum-likelihood phylogenetic trees from multiple sequence alignments of concatenated core SNVs. Reference genomes were selected based on the clinical laboratory MALDI-TOF classification of BSI isolates. Phylogenetic trees demonstrate a higher level of intra-patient versus inter-patient genetic relatedness among colonizing and infecting organisms (Figure 4). BSI and intestinal strains of typically enteric species such as *Enterococcus faecium* (patient 25), *Klebsiella pneumoniae* (patient 2), *E. coli* (patient 3, patient 7), and *S. mitis* (patient 22) are closely phylogenetically related. Unexpectedly, we observe close intra-patient strain relatedness between gut and BSI for typically non-enteric taxa including *S. epidermidis* strains (patient 13) and *Pseudomonas aeruginosa* (patient 19, not pictured). Phylogenetic trees for *P. aeruginosa* and *Enterobacter cloacae* are not shown as these species were not observed in any other samples.

**Figure 4.**
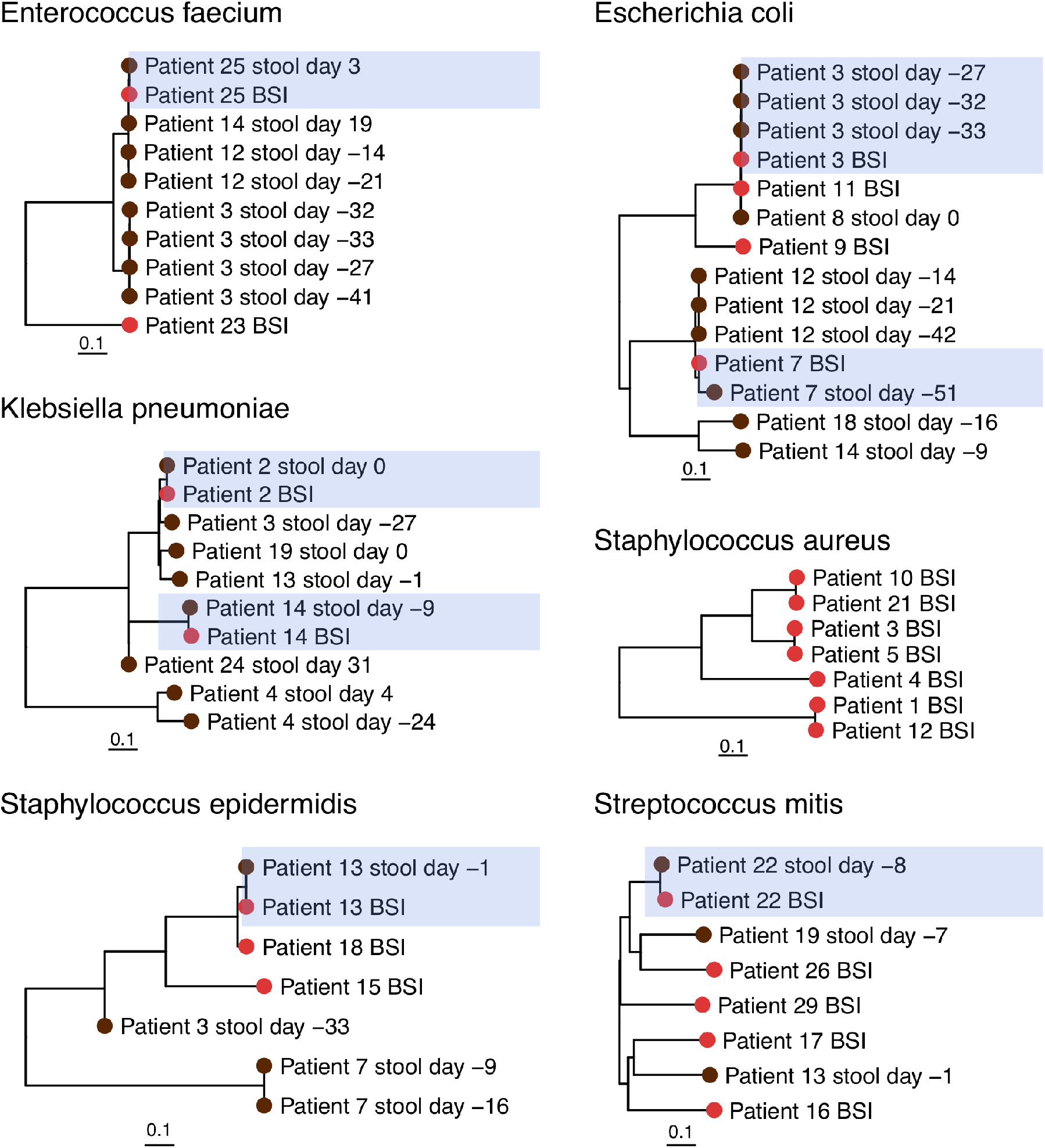
Intra-patient samples are more closely related than inter-patient samples. Branch tip colors indicate stool and BSI samples from the same patient. Samples from the same patient are more closely phylogenetically related than other samples.

### Single-nucleotide variants reveal strain identity

We next used StrainSifter to compare single-nucleotide variants (SNVs) between strains in our sample collection. We executed the StrainSifter pipeline for each draft genome, with all stool short read datasets as input. StrainSifter identified samples with sufficient coverage (at least 5X coverage across 50% or more of the genome) of the target organism in the gut metagenome and determined the number of variants shared between each pair of input samples that contained the target strain.

We observe 1 SNV between the BSI and gut content of Patient 13 who developed a *S. epidermidis* BSI (Table 3). This indicates that the bloodstream strain is concordant to the strain found in the gut 1 day prior. Patient 3 developed an *E. coli* bloodstream infection and we observe zero SNVs between the BSI strain and all samples 33, 32, and 27 days prior to the BSI, indicating that the identical E. coli BSI strain colonized the gut over a month before the onset of infection. Further, we observe zero discriminating SNVs between identical strains of *Pseudomonas aeruginosa* in both the blood and stool specimens. Notably, all stool samples containing the BSI-causing organism at sufficient coverage to profile with StrainSifter were more closely related to their respective BSI isolate based on SNV distance than to any sample from other patients. None of the 30 patients included in our study had sufficient *Staphylococcus aureus* in their stool samples to profile with StrainSifter, indicating that this organism infrequently colonizes the gut of HCT patients, even at low abundance.

**Table 3.**

| BSI organism | Sample 1 | Sample 2 | SNVs | Bases compared | SNVs per base compared | Total bases | Percentage of bases compared |
| --- | --- | --- | --- | --- | --- | --- | --- |
| Enterobacter cloacae | Patient 2 BSI | Patient 2 stool day 0 | 5 | 4499921 | 1.11E-06 | 4552154 | 98.85% |
| Enterococcus faecium | Patient 25 BSI | Patient 25 stool day 3 | 2 | 2931442 | 6.82E-07 | 2935264 | 99.87% |
| Escherichia coli | Patient 3 BSI | Patient 3 stool day -33 | 0 | 5032224 | 0.00E+00 | 5056158 | 99.53% |
| Escherichia coli | Patient 3 BSI | Patient 3 stool day -32 | 0 | 5032230 | 0 | 5056158 | 99.53% |
| Escherichia coli | Patient 3 BSI | Patient 3 stool day -27 | 0 | 5030454 | 0 | 5056158 | 99.49% |
| Escherichia coli | Patient 7 BSI | Patient 7 stool day -51 | 259 | 4388254 | 5.90E-05 | 5267890 | 83.30% |
| Klebsiella pneumoniae | Patient 2 BSI | Patient 2 stool day 0 | 4 | 5379270 | 7.44E-07 | 5390737 | 99.79% |
| Klebsiella pneumoniae | Patient 14<br>BSI | Patient 14 stool day -9 | 72 | 3924126 | 1.83E-05 | 5407718 | 72.57% |
| Pseudomonas aeruginosa | Patient 19<br>BSI | Patient 19 stool day 0 | 0 | 6830248 | 0 | 6882743 | 99.24% |
| Staphylococcus<br>epidermidis | Patient 13<br>BSI | Patient 13 stool day -1 | 1 | 2592411 | 3.86E-07 | 2604609 | 99.53% |
| Streptococcus mitis | Patient 22<br>BSI | Patient 22 stool day -8 | 28 | 1345797 | 2.08E-05 | 2084062 | 64.58% |

### *In silico* multilocus sequence typing is concordant with StrainSifter results

To compare whole-genome sequencing-based approaches to traditional strain typing methods, we performed *in silico* multilocus sequence typing (MLST) on BSI genomes and assembled stool metagenomes (Table 3). In many instances, MLST types for gut metagenome strains could not be resolved or could only be partially detected, likely due to low abundance or absence of the target organism in the gut community. MLST data is concordant with StrainSifter SNV profiling in only 4 infections for which MLST schemes exist for the bloodstream isolate of interest (Table S7). MLST types could not be determined for either of patient 2’s BSI isolates *(K. pneumoniae* and *E. cloacae)* because the sequences at several loci in each genome were not present in the respective MLST databases. We were unable to type *Rothia mucilaginosa* and *Streptococcus mitis,* as standard MLST schemes do not exist for these organisms.

## Discussion

### StrainSifter is a useful pipeline for strain detection within metagenomes

High resolution strain comparisons are required to identify the origins of bloodstream infections using metagenomic samples. While many bioinformatic tools exist for strain comparison between clinical isolates or SNV profiling between metagenomes, no pipeline exists for the specific purpose of detecting the presence of a single strain in a metagenome. In this study, we have developed StrainSifter, a straightforward and facile pipeline for identifying phylogenetic relationships between isolate and metagenomic WGS samples.

In our study, StrainSifter was able to precisely and comprehensively compare gut metagenomic content to the genomes of diverse bloodstream isolates using high resolution WGS. Although techniques such as *in silico* MLST are able to compare genomes to metagenomes, MLST loci are only 450 to 500 base pairs on average, and standard 7-gene MLST schemes therefore analyze less than 4 kilobases of DNA - on the order of 0.1% of the average bacterial genome. By examining genome-wide variants, StrainSifter greatly increases the confidence of strain comparisons.

Using StrainSifter, we found that the enteric organisms *E. faecium, E.coli, K. pneumoniae, E. cloacae,* and *S. mitis* were concordant between blood and stool microbiota samples. This confirms what has previously been reported in the literature (Tancrede and Andremont 1985; Samet et al. 2013). Previously published studies have indicated that gut domination with a pathogen is a risk factor for subsequent bloodstream infection (Taur et al. 2012; Ubeda et al. 2010; “Centers for Disease Control and Prevention” 2018). While *E. coli* and *E. faecium* domination events were observed preceding BSI, we find *K. pneumoniae*, *E. cloacae,* and *S. epidermidis* in relatively low abundance in stool samples on the day of BSI and one day prior.

Interestingly, we also find that typically non-enteric organisms *S. epidermidis* and *P. aeruginosa* were not only present in the gut, but strains within the gut microbiota and bloodstream were concordant. Gut colonization by *S. epidermidis* and *P. aeruginosa* is unexpected given current National Health and Safety guidelines delimiting enteric versus non-enteric organisms (Taur et al. 2012; Ubeda et al. 2010; “Centers for Disease Control and Prevention” 2018; Advani et al. 2017). Yet, colonization by atypical organisms may not be surprising given the degree of disruption of the intestinal microbiota by antibiotics, chemotherapy, and radiation. While *P. aeruginosa* intestinal colonization has been previously documented (Nesher et al. 2014), intestinal colonization with coagulase-negative *Staphylococcus* is not well described. In our own study, multiple stool samples were determined to have more than 80% relative abundance of *S. epidermidis*. In addition, we observe the same colonizing strain of coagulase-negative *Staphylococcus* over time in the same individual (patient 7), arguing against only transient membership within the intestinal microbiome.

Previous research using existing strain typing and culture-based strategies has only tentatively demonstrated evidence for gut translocation of coagulase-negative *Staphylococcus spp.* and *P. aeruginosa* in immunocompromised patients (Wade et al. 1982; Rotstein et al. 1988; Nesher et al. 2014; MacFie et al. 1999; Costa et al. 2006). Here our straightforward approach allows for greater precision and confidence in strain comparisons between the gut microbiota and infecting organism in the bloodstream. Given that our analysis focused entirely on the stool microbiota, we cannot rule out the possibility of the same strain of coagulase-negative *Staphylococcus* colonizing the entire body surface--including the gut, nares, skin, or central line--from which the infection may have originated instead. Previous research has suggested, however, that the majority of patients with coagulase-negative *Staphylococcus* bloodstream infection are colonized with a multitude of strains that differ between the intestine and the skin (Herwaldt et al. 1992).

Determining whether two strains are related based on the number of SNVs that differ between their genomes is not straightforward. While approximate bacterial mutation rates due to genetic drift have been established, most estimates are derived from experiments in culture media. As gut and bloodstream environments impose many selective pressures beyond genetic drift, such estimates are likely not relevant in the clinical context. For instance, it is possible that bacterial pathogens that originate from the intestine and translocate into the bloodstream may acquire single nucleotide variants over time due to selective pressures. For this reason, it is difficult to define a threshold number of SNVs beyond which two strains are not considered to be the same. Regardless of the absolute number of SNVs between strains or across all strains analyzed with StrainSifter, strains from the same patient consistently share fewer SNVs compared to those from other patients.

Although StrainSifter can precisely identify shared variants between genomes and metagenomes, it is limited to profiling only the dominant strain of a given organism in a community. However, it has been shown that gut metagenomes frequently contain only one predominant strain of each species, so StrainSifter is likely to function optimally in most situations (Truong et al. 2017). As is the case with existing tools for metagenomic strain comparison, StrainSifter is unable to analyze low-abundance strains since adequate coverage (5X) is required for confident variant calling.

### Future directions

StrainSifter can be used to precisely identify strains within relatively biomass-rich metagenomic samples without relying on external reference genome databases in which representation of opportunistic pathogens may be lacking. Compared to stool, samples from the nares, skin, or the environment typically have far less microbial biomass, limiting the number of sequencing reads that can be obtained. In order to provide the most precise epidemiologic information about the origins of bloodstream and other infections, StrainSifter will need to be tested in low microbial biomass samples in a larger prospective study of immunocompromised patients. Given the high resolution of WGS-based strain comparisons, we may indeed discover instances of organisms thought to originate from non-enteric sites instead colonizing and translocating from the intestine in severely immunocompromised patients, as was potentially demonstrated by *S. epidermidis* and *P. aeruginosa* in our study. Given the growing body of research on therapies to improve gut microbiota diversity and bolster colonization resistance against pathogens, more precisely identifying the origins of bloodstream infections in the stem cell transplant population may influence how hospitals and healthcare providers can most effectively work to prevent infections (Galloway-Peña, Jenq, and Shelburne 2017). Finally, once StrainSifter is able to establish the origins of an infection, the wealth of information offered by WGS can be interrogated to identify potential functional differences, including in gene content and metabolic pathways, between strains colonizing the gut, nares, or skin that result in disseminated infections compared to those that do not.

In conclusion, we have shown that StrainSifter offers a straightforward bioinformatic pipeline to precisely compare bacterial isolates to metagenomic samples, circumventing the need for external reference genome databases or culture-based methods of strain comparison. We successfully applied StrainSifter to blood culture isolates and stool samples from hematopoietic stem cell transplant patients and found classically non-enteric bacteria in the gut microbiome that were concordant with bloodstream isolates. We anticipate that StrainSifter will be an easy-to-use tool in the context of hospital or infectious outbreak epidemiologic studies, and that characterization of the microbiome dynamics that occur prior to infection may help us precisely identify potential reservoirs of pathogens.

## Acknowledgements

We thank Joyce Kang for her assistance with stool sample processing, as well the other members of the Bhatt Lab for providing feedback on the study design, bioinformatics pipeline, and manuscript revisions. We would like to thank Nick Greenfield and the One Codex team for help with using their platform. We appreciate Drs. Matthew Kelly, Chris Severyn, and Doyle Ward for their feedback and ideas on the manuscript. We would especially like to thank the patients and nurses on the Blood and Marrow Transplant service for their enthusiastic participation in this project. This work was supported in part by National Science Foundation Graduate Research Fellowship (FBT), the National Institutes of Health, National Center for Advancing Translational Science, Clinical and Translational Science Award KL2 TR000083 and UL1 TR001085 (TMA). ASB was funded in part by the National Cancer Institute NIH K08 award, #CA184420, Damon Runyon Clinical Investigator Award, and Amy Strelzer Manasevit Award. The content is solely the responsibility of the authors and does not necessarily represent the official views of the NIH.

